# Development of a locus-specific HRM assay for DNA methylation analysis of the SHANK3 gene

**DOI:** 10.1101/2025.09.03.673955

**Authors:** Sema Tiryaki

**Affiliations:** TUBITAK National Metrology Institute (TUBITAK UME), Chemistry Group, Bioanalysis Laboratory, PK 54, 41470 Kocaeli, Türkiye

**Keywords:** SHANK3, DNA methylation, CpG island, high-resolution melting, bisulfite conversion, neuropsychiatric disorders

## Abstract

DNA methylation within CpG islands is a key epigenetic mechanism regulating gene expression. SHANK3 encodes a synaptic scaffolding protein essential for neurodevelopment and synaptic function, and aberrant SHANK3 methylation has been implicated in neuropsychiatric disorders. To enable reliable locus-specific investigation of SHANK3 epigenetic regulation, we developed a high-resolution melting (HRM) assay. In silico screening identified a CpG-rich region upstream of exon 3 as the most suitable locus for assay design. Bisulfite-converted sequences representing fully methylated and unmethylated states were used to generate three primer sets, of which two successfully amplified the target region and produced distinct melting profiles discriminating methylated from unmethylated templates. The assay was optimized on two HRM platforms (LightCycler® 480 and CFX96), and conversion efficiency was confirmed with commercial control DNAs. This locus-specific HRM assay provides a methodological framework for qualitative SHANK3 methylation analysis and represents a promising tool for future validation studies and potential clinical investigations in neurodevelopmental and neuropsychiatric disorders.

## Introduction

DNA methylation is one of the most studied epigenetic modifications and plays a critical role in gene regulation, chromatin remodeling, and cellular differentiation. Aberrant methylation, particularly in promoter regions, has been consistently associated with transcriptional silencing of tumor suppressor genes and the activation of oncogenic pathways[1–3]. Promoter hypermethylation is one of the earliest events in carcinogenesis, providing unique opportunities for its use as a biomarker in both diagnostic and prognostic contexts [4–6]. Due to its chemical stability, cancer-type specificity, and translational potential, promoter methylation has emerged as one of the most powerful biomarker candidates in precision medicine[7–10].

Although most studies have traditionally focused on promoter regions upstream of the first exon, methylation analysis is not restricted to these loci. Several studies have reported methylation in the first exon, internal exons, and intragenic CpG islands, highlighting that exonic methylation may influence transcription regulation, alternative splicing, and gene silencing [11–14] Thus, the investigation of methylation across exonic regions represents a growing area of research with potential diagnostic relevance.

One gene of particular interest in this context is SHANK3, a postsynaptic density protein-coding gene essential for synaptic organization and function. SHANK3 mutations and epigenetic alterations have been extensively studied in autism spectrum disorders (ASD), neurodevelopmental disorders (NDDs), and synaptopathies such as Phelan-McDermid syndrome (PMS) [15–17]. Gene body methylation of SHANK3 has been linked to downregulation of mRNA expression in animal models [16], while tissue-specific methylation changes in intragenic/promoter-proximal CpG islands have been observed in ASD brain tissues. Furthermore, SHANK3 promoter methylation has been proposed as a biomarker in a subset of studies investigating neuronal tissue methylation patterns and environmental exposures [15,17].

Although the available evidence is still limited, SHANK3-related genetic and epigenetic changes—including mutations, gene body methylation, and promoter methylation—are increasingly recognized for their diagnostic and therapeutic potential. Investigating SHANK3 methylation across both promoter and exonic regions represents a promising avenue for biomarker discovery and for elucidating disease mechanisms in ASD, PMS, and schizophrenia. In this study, rather than restricting our analysis to the commonly investigated promoter region upstream of exon 1—which we initially examined but found unsuitable for CpG island definition and primer design—we focused on a CpG-rich island located between exons 2 and 3 of the SHANK3 gene. Our assays were designed to specifically target this exonic region, thereby extending SHANK3 methylation analysis beyond the traditionally examined promoter. This work provides a methodological framework that may support future clinical and translational investigations of SHANK3 methylation.

## Material and Method

### Selection of Target Region by in silico CpG Island Screening and Primer Design

The SHANK3 gene sequence (NCBI Gene ID: 85358) was examined to identify CpG islands suitable for promoter methylation analysis by HRM. CpG Island Grapher (Hahn Lab utilities, http://canna.cau.ac.kr/util/) was used, with CpG islands defined as regions with GC content >55–60 and a CpG observed/expected (O/E) ratio >0.6. A CpG island fulfilling these thresholds (Figure 1) was identified upstream of exon 3, and a 1,135 bp fragment spanning this region was selected for assay design.

**Figure 1.**
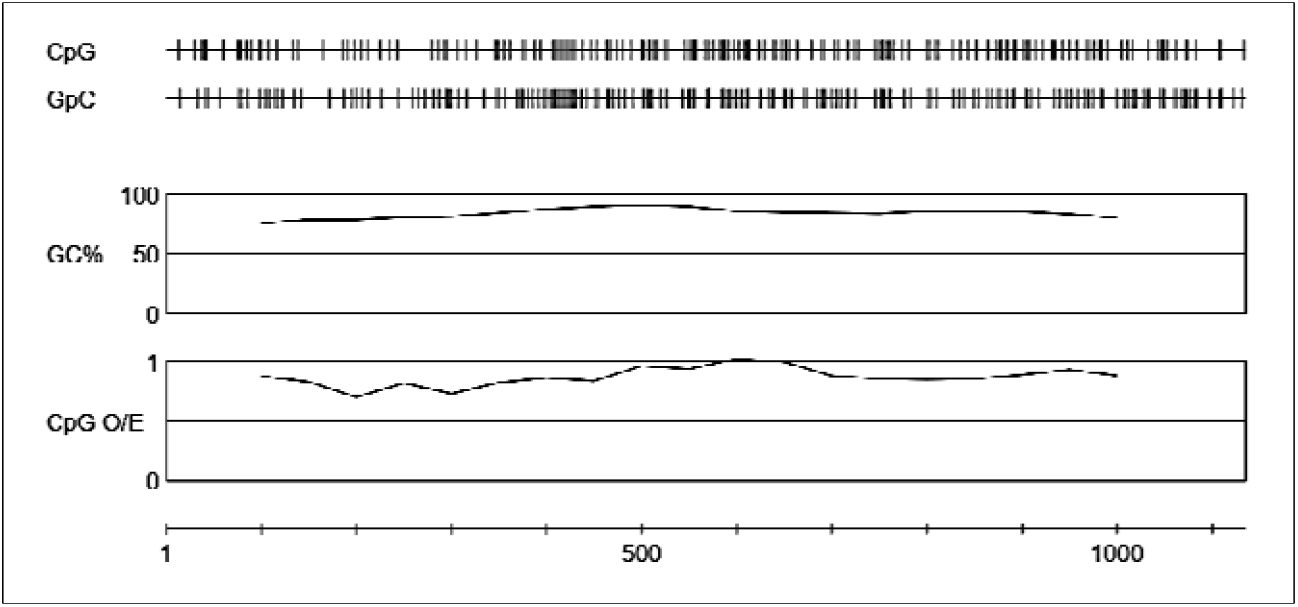
1000kb upstream Exon 3 CpG island screeening.

The “Bisulfite Conversion” tool (Hahn Lab utilities) used to obtain bisulfite-converted (BC) sequences for fully methylated (M) and fully unmethylated (UnM) DNA. The original, M-BC and UnM-BC sequences were aligned (Supplementary Figure S1) Primers were designed to target regions that remain unchanged after bisulfite conversion, with the following considerations: minimization of cytosines (if unavoidable, only one C and not positioned at the 3′ end), amplicon size between 80–300 bp, and avoidance of homopolymers, repeats, and 3′ complementarity. Three primer pairs (Primer Sets 1–3) were generated. Candidate primers were evaluated using IDT OligoAnalyzer to assess secondary structures and hairpin formation, and to calculate melting temperatures (Tm). Predicted melting temperatures of the expected amplicons were further determined using Amplicon3Plus (https://www.primer3plus.com/amplicon3plus.html). Oligonucleotides were synthesized commercially (Oligomer Biyoteknoloji, Ankara, Türkiye). The primer sequences are summarized in Table 1. Detailed information on amplicon sequences, amplicon length, and predicted melting temperatures for methylated and unmethylated templates is provided in Supplementary Table S1-S2.

**Table 1.**
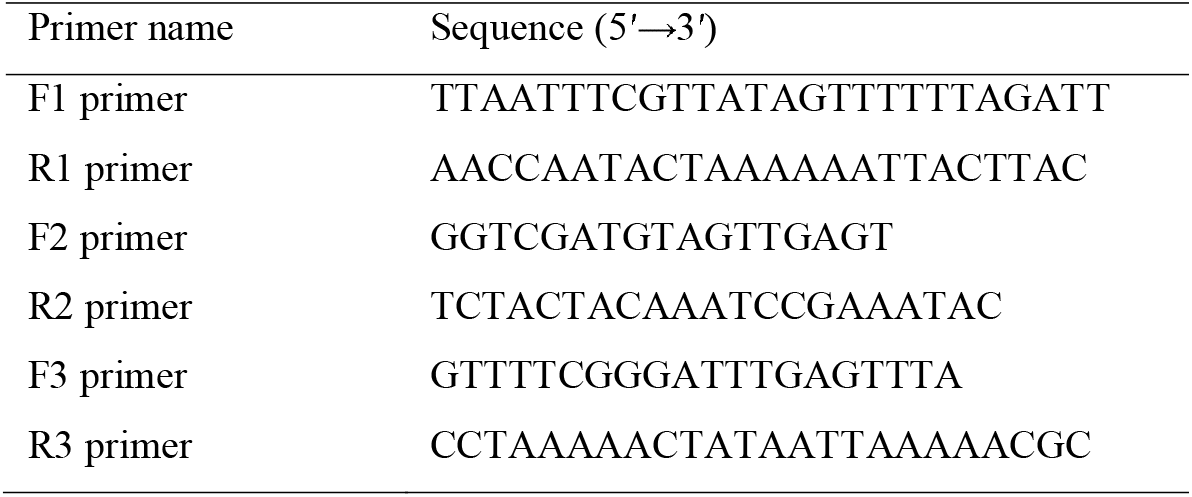
Primer sequences designed for SHANK3 methylation analysis.

### Bisulphite Conversion of gDNAs

For bisulfite conversion, the EZ DNA Methylation™ Kit (Zymo Research) was used following the manufacturer’s protocol. The Human Methylated & Non-methylated DNA Set (D5014, Zymo Research), consisting of paired control DNAs (methylated and unmethylated, 250 ng/µL), was included to monitor conversion efficiency and to serve as PCR controls. Prior to bisulfite treatment, control DNAs were diluted 1:10, and 4 µL of each was subjected to conversion. Converted DNA was eluted in 10 µL and subsequently diluted 1:5 prior to PCR/HRM analysis.

### qPCR and HRM Analysis

All assays were carried out on a LightCycler® 480 System (Roche) and a CFX96 Real-Time PCR Detection System (Bio-Rad). PCR reactions were performed in a total volume of 20 µL, containing SOLIS BIODYNE Hot FirePol® EvaGreen® HRM Mix (5×), 400 nM of each primer, and 4 µL of bisulfite-converted DNA template. Post-conversion DNA was not quantified; instead, a fixed template volume (4 µL per 20 µL reaction) was used. Annealing temperature optimization was performed using a gradient PCR (52–60 °C) with converted methylated and unmethylated control DNAs.

HRM analyses were performed using the LightCycler® 480 Software (Gene Scanning module, Roche) and CFX Maestro Software with the Precision Melt Analysis module (Bio-Rad). In all experiments, reactions were run in duplicate using bisulfite-converted control gDNAs (methylated and unmethylated) as templates. No-template controls (NTCs) were consistently included in parallel to monitor nonspecific amplification.

## Results

### Bisulfite conversion controls

Bisulfite conversion was successfully verified using the DAPK1 control assay included in the commercial kit (Zymo D5014). Amplification was obtained at the expected melting temperatures, confirming conversion of unmethylated cytosines and overall PCR competency.

### Amplification and melt curve evaluation of SHANK3 primer sets

Among the three primer sets tested, Primer Sets 1 and 3 produced amplifications, yielding melt peaks that corresponded to the predicted melting temperatures of the amplicons (Supplementary Table S2; Figure 2, Figure 3). Primer Set 2, however, showed weak or inconsistent amplification and was therefore excluded from further analyses. In no-template controls (NTCs), non-specific low-temperature peaks (likely primer-dimer products) were occasionally observed below 70 °C; these were absent in reactions containing template DNA and did not interfere with interpretation of the specific amplicon peaks (see Supplementary Figure S1).

**Figure 2.**
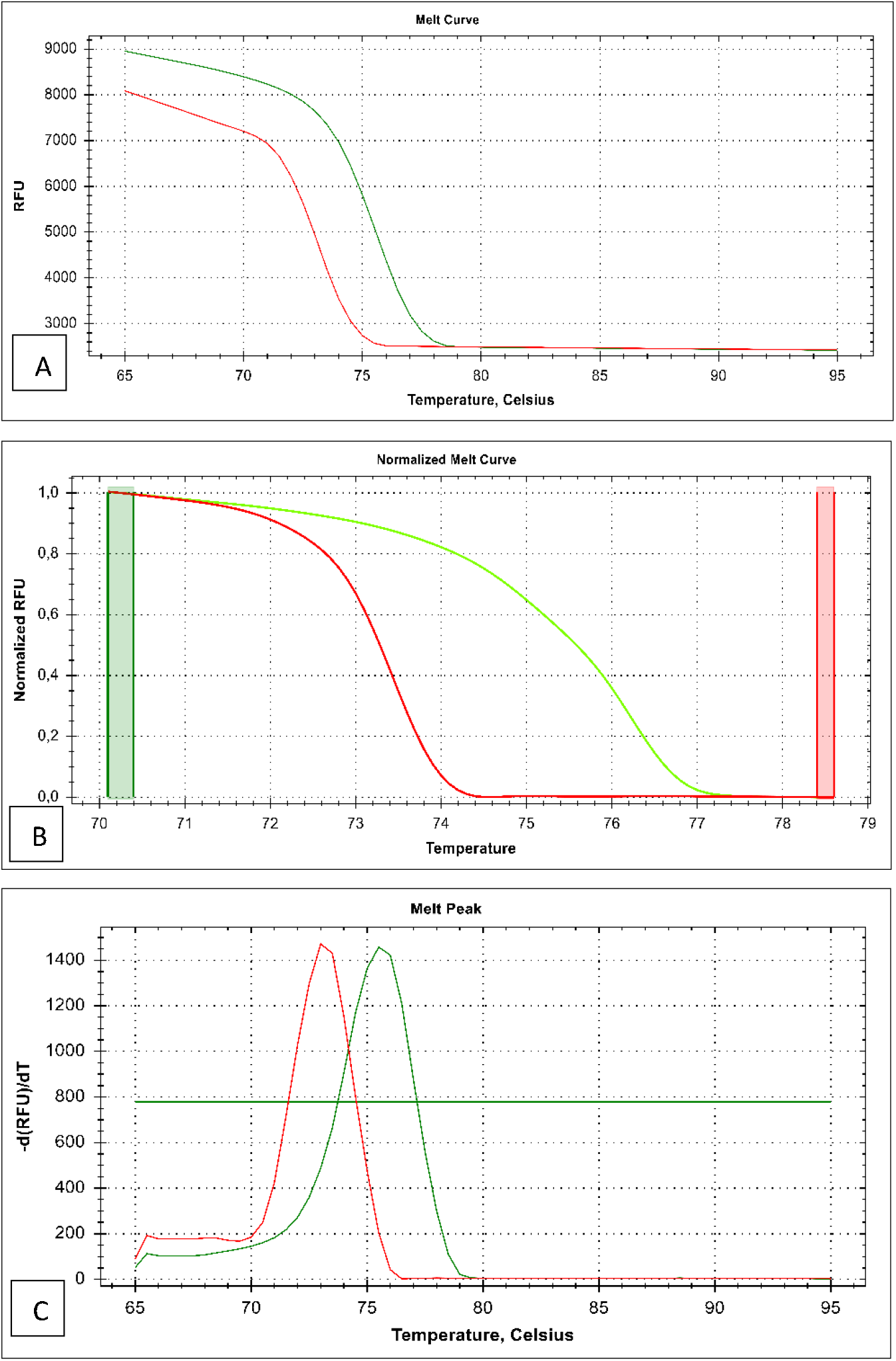
Amplification and melt curve evaluation of SHANK3 Primer set 1. (A) Representative amplification curves obtained with bisulfite-converted methylated (green) and unmethylated (red) control DNAs using Primer Set 1. (B) Normalized melt curves showing distinct separation between methylated and unmethylated templates. (C) Corresponding derivative melt peaks with Tm values of 73 °C and 75,5 °C for unmethylated and methylated templates, respectively.

**Figure 3.**
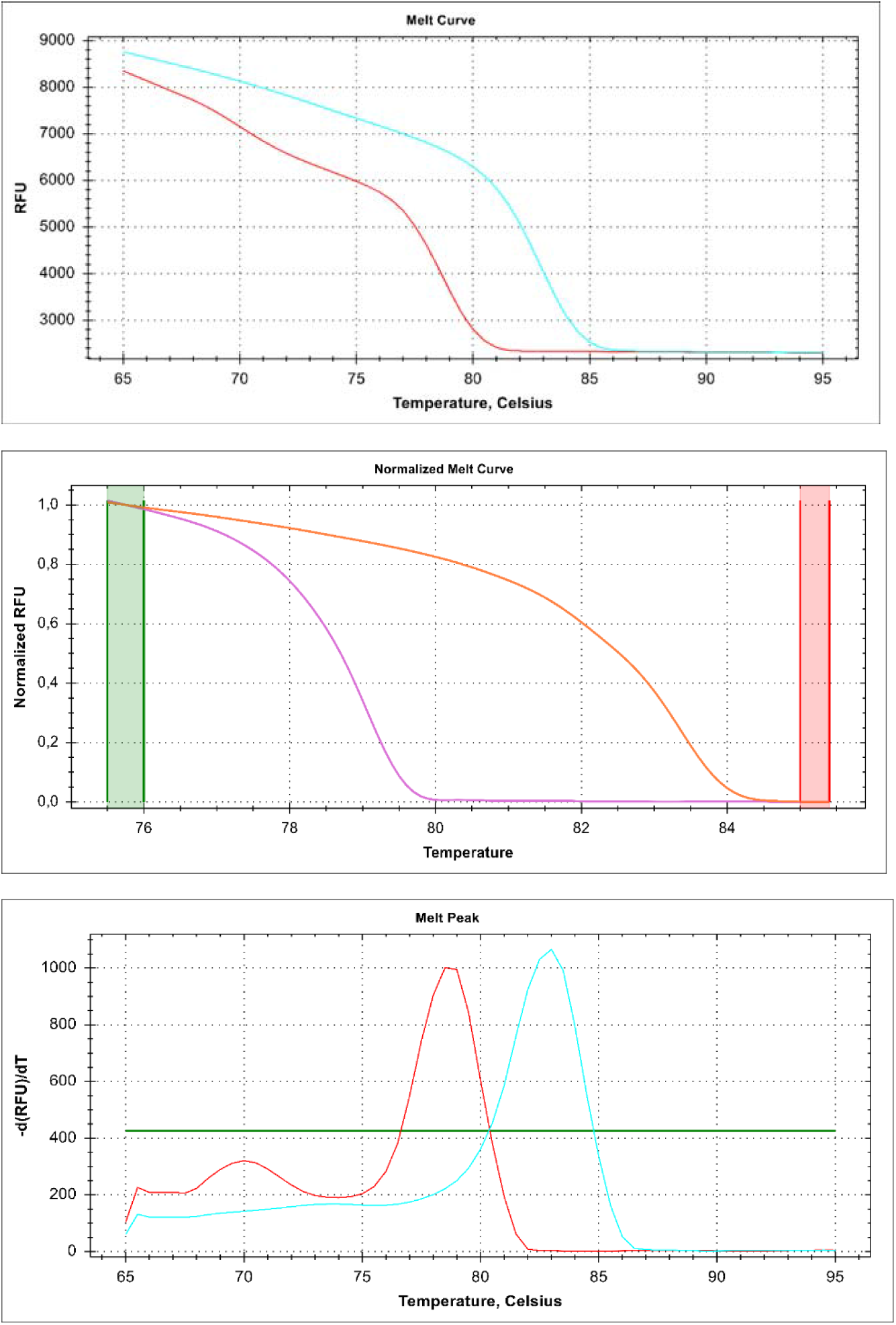
Amplification and melt curve evaluation of SHANK3 Primer set 3. (A) Representative amplification curves obtained with bisulfite-converted methylated (green) and unmethylated (red) control DNAs using Primer Set 3. (B) Normalized melt curves showing distinct separation between methylated and unmethylated templates. (C) Corresponding derivative melt peaks with Tm values of 78,5 °C and 83 °C for unmethylated and methylated templates, respectively.

### Gradient PCR optimization

Gradient PCR (52–60 °C) was carried out to determine optimal annealing conditions. For both Primer Sets 1 and 3, an annealing temperature of 54 °C yielded the most efficient and specific amplification, as evidenced by sharp melt peaks with minimal background signal.

### Non-template controls and nonspecific amplification

In Primer Set 3, no-template controls (NTCs) showed late-cycle amplification. The derivative melting curves indicated that these non-specific products had melting temperatures lower than the target amplicon (Figure 3) and therefore did not interfere with data interpretation. The presence of the correct product was still readily confirmed in template-containing wells.

### Assay applicability

Primer Set 1 and 3 enabled discriminations between methylated and unmethylated control DNAs (Figure 2, Figure 3), supporting their use for SHANK3 methylation analysis by HRM. However, based on current data, the assay is most appropriate for qualitative “methylated vs. unmethylated” classification. Further validation, including determination of the limit of detection (LOD), limit of quantification (LOQ), and analytical sensitivity parameters, would be required to establish quantitative performance.

## Discussion

In this study, a high-resolution melting (HRM) assay was developed and tested for the analysis of DNA methylation within a CpG island located upstream of exon 3 of the SHANK3 gene. By integrating in silico CpG island prediction, bisulfite conversion, and HRM-based analysis, we established a locus-specific workflow that enabled the discrimination of methylated and unmethylated templates.

The bisulfite conversion process was verified using the DAPK1 control assay and commercially available methylated and unmethylated DNAs. The successful amplification of these controls at expected melting temperatures confirmed the efficiency of conversion and the validity of subsequent HRM analysis. Among the three primer sets designed, Primer Sets 1 and 3 demonstrated consistent amplification and reliable discrimination between methylated and unmethylated templates, while Set 2 was less efficient. Gradient PCR optimization further established 54 °C as the optimal annealing temperature, ensuring reproducible performance.

Some nonspecific amplification events were observed in non-template controls (NTCs) and, to a lesser extent, in template-containing wells. However, these products displayed melting temperatures distinct from the expected amplicon, and thus did not interfere with interpretation. This observation underscores the importance of HRM curve analysis as a powerful tool for distinguishing true targets from nonspecific products.

The established assay is particularly suitable for qualitative “methylated versus unmethylated” classification. At this stage, the method cannot be considered quantitative, as validation parameters such as the limit of detection (LOD), limit of quantification (LOQ), and analytical sensitivity were not determined. Future work should address these parameters and evaluate assay performance across independent DNA samples to fully establish analytical robustness.

Despite these limitations, the present work provides an essential methodological foundation for SHANK3 methylation analysis. The primer sets and optimized HRM conditions reported here can serve as a basis for further investigations of SHANK3 epigenetic regulation in both research and diagnostic contexts.

## Conclusion

This study describes the development of a locus-specific HRM assay for the analysis of DNA methylation in the SHANK3 gene. By combining in silico CpG island prediction, bisulfite conversion, and HRM optimization, we established primer sets and experimental conditions that enable clear discrimination between methylated and unmethylated templates. While the current assay is best suited for qualitative methylation assessment, further validation including LOD, LOQ, and analytical sensitivity studies will be required to extend its application to quantitative analyses. Overall, this work provides a methodological framework that may support future epigenetic investigations of SHANK3.

## Acknowledgments

The author gratefully acknowledges the Bioanalysis Laboratory at TÜBİTAK UME for providing the facilities and resources that made this work possible.

## Data availability statement

All data generated or analyzed during this study are included in this published article and its supplementary information files.

## References

[1] Geissler F, Nesic K, Kondrashova O, et al. The role of aberrant DNA methylation in cancer initiation and clinical impacts. Ther Adv Med Oncol 2024;16:17588359231220512. 10.1177/17588359231220511.

[2] Wajed SA, Laird PW, DeMeester TR. DNA Methylation: An Alternative Pathway to Cancer. Ann Surg 2001;234:10. 10.1097/00000658-200107000-00003.

[3] Wu X, Zhang Y. TET-mediated active DNA demethylation: Mechanism, function and beyond. Nat Rev Genet 2017;18:517–34. 10.1038/NRG.2017.33;SUBJMETA.

[4] Leung WK, To KF, Chu ESH, et al. Potential diagnostic and prognostic values of detecting promoter hypermethylation in the serum of patients with gastric cancer. Br J Cancer 2005;92:2190–4. 10.1038/SJ.BJC.6602636.

[5] Lavoro A, Ricci D, Gattuso G, et al. Recent advances on gene-related DNA methylation in cancer diagnosis, prognosis, and treatment: a clinical perspective. Clin Epigenetics 2025;17. 10.1186/S13148-025-01884-2.

[6] Müller D, Győrffy B. DNA methylation-based diagnostic, prognostic, and predictive biomarkers in colorectal cancer. Biochim Biophys Acta Rev Cancer 2022;1877. 10.1016/j.bbcan.2022.188722.

[7] Locke WJ, Guanzon D, Ma C, et al. DNA Methylation Cancer Biomarkers: Translation to the Clinic. Front Genet 2019;10:1150. 10.3389/FGENE.2019.01150.

[8] Mikeska T, Craig JM. DNA Methylation Biomarkers: Cancer and Beyond. Genes (Basel) 2014;5:821. 10.3390/GENES5030821.

[9] Lavoro A, Ricci D, Gattuso G, et al. Recent advances on gene-related DNA methylation in cancer diagnosis, prognosis, and treatment: a clinical perspective. Clin Epigenetics 2025;17:1–42. 10.1186/S13148-025-01884-2/TABLES/3.

[10] Mikeska T, Bock C, Do H, et al. DNA methylation biomarkers in cancer: progress towards clinical implementation. Expert Rev Mol Diagn 2012;12:473–87. 10.1586/ERM.12.45.

[11] Brenet F, Moh M, Funk P, et al. DNA Methylation of the First Exon Is Tightly Linked to Transcriptional Silencing. PLoS One 2011;6:e14524. 10.1371/JOURNAL.PONE.0014524.

[12] Li S, Zhang J, Huang S, et al. Genome-wide analysis reveals that exon methylation facilitates its selective usage in the human transcriptome. Brief Bioinform 2018;19:754–64. 10.1093/BIB/BBX019.

[13] Maunakea AK, Chepelev I, Cui K, et al. Intragenic DNA methylation modulates alternative splicing by recruiting MeCP2 to promote exon recognition. Cell Res 2013;23:1256–69. 10.1038/CR.2013.110.

[14] Gelfman S, Cohen N, Yearim A, et al. DNA-methylation effect on cotranscriptional splicing is dependent on GC architecture of the exon–intron structure. Genome Res 2013;23:789. 10.1101/GR.143503.112.

[15] Zhu L, Wang X, Li XL, et al. Epigenetic dysregulation of SHANK3 in brain tissues from individuals with autism spectrum disorders. Hum Mol Genet 2014;23:1563–78. 10.1093/HMG/DDT547.

[16] Barua S, Kuizon S, Chadman KK, et al. Single-base resolution of mouse offspring brain methylome reveals epigenome modifications caused by gestational folic acid. Epigenetics Chromatin 2014;7:1–15. 10.1186/1756-8935-7-3/FIGURES/3.

[17] Zhou Q, Tian Y, Xu C, et al. Prenatal and postnatal traffic pollution exposure, DNA methylation in Shank3 and MeCP2 promoter regions, H3K4me3 and H3K27me3 and sociability in rats’ offspring. Clin Epigenetics 2021;13. 10.1186/S13148-021-01170-X.

