## Supplementary Figures for "Development of a locus-specific HRM assay for DNA methylation analysis of the SHANK3 gene"

> Supplementary Figure S1 / SHANK3- Upstream of Exon 3- 'F Primer : R Primer:

RefSeq Row 1 : tctttccccttctCGccctaccttggtcttCGtgcccCGaCGCGgcccccaccccagctcCGggaccccttcctcctcccccccaacccCGtcacagcccccagaccccCGcccCGtggctCGgccccCGccctcCGcacacacctcc

BC-methylated Row 1 : TTTTTTTTTTTTTCGTTTTATTTTGGTTTTCGTGTTTCGACGCGGTTTTTATTTTAGTTTCGGGATTTTTTTTTTTTTTTTTTTTTAATTTCGTTATAGTTTTTTAGATTTTCGTTTCGTGGTTCGGTTTTCGTTTTTCGTATATATTTTT

BC-unmethylated Row 1 : TTTTTTTTTTTTTTGTTTTATTTTGGTTTTTGTGTTTTGATGTGGTTTTTATTTTAGTTTTGGGATTTTTTTTTTTTTTTTTTTTTAATTTTGTTATAGTTTTTTAGATTTTTGTTTTGTGGTTTGGTTTTTGTTTTTTGTATATATTTTT

RefSeq Row 2 : CgcccccaccCGggacccCGcaagtaaccccccagcactggccctgagccctccCGgccccCGcctcCGgCGcagccccctCGccaccccCGcttccctccCGtctcaggccccctccccCGcCGccccCGccccCGgggaaggcaggC

BC-methylated Row 2 :CGTTTTTATTCGGGATTTCGTAAGTAATTTTTTAGTATTGGT*TTTGAGTTTTTTCGGTTTTCGTTTTCGGCGTAGTTTTTTCGTTATTTTCGTTTTTTTTTCGTTTTAGGTTTTTTTTTTTCGTCGTTTTCGTTTTCGGGGAAGGTAGGC*

BC-unmethylated Row 2 :TGTTTTTATTTGGGATTTTGTAAGTAATTTTTTAGTATTGGTTTTGAGTTTTTTGGTTTTTGTTTTTGGTGTAGTTTTTTGTTATTTTGTTTTTTTTTTGTTTTAGGTTTTTTTTTTTGTTGTTTTTGTTTTTGGGGAAGGTAGGT

RefSeq Row 3 : GcCGagctgagcCGgggcCGatgcagctgagcCGCGcCGcCGcCGcCGcCGcCGccccctgCGgagccccCGgagcCGctgtccccCGCGcCGgcccCGgcccCGgccccccCGgcccccctccCGCGcagCGCGgcCGaCGgggctcC

BC-methylated Row 3 : *GTCGAGTTGAGTCGGGGTCGATGTAGTTGAGT*CGCGTCGTCGTCGTCGTCGTCGTCGTTTTTGCGGAGTTTTCGGAGTCGTTGTTTTTCGCGTCGGTTTCGGTTTCGGTTTTTTCGGTTTTTTTTCGCGTAGCGCGGTCGACGGGGTTTC

BC-unmethylated Row 3 :GTTGAGTTGAGTTGGGGTTGATGTAGTTGAGTTGTGTTGTTGTTGTTGTTGTTGTTGTTTTTGTGGAGTTTTTGGAGTTGTTGTTTTTTGTGTTGGTTTTGGTTTTGGTTTTTTTGGTTTTTTTTTGTGTAGTGTGGTTGATGGGGTTTT

RefSeq Row 4 : GgCGggggggaagggggggcCGgggCGcCGCGCGgagtcccCGggCGctcCGttcccCGgCGCGagCGgcccCGgccCGggccCGgCGCGgggatggaCGgcccCGgggccagCGcCGtggtCGtgCGCGtCGgcatccCGgacctgcagcaga

BC-methylated Row 4 : GGCGGGGGGGAAGGGGGGGTCGGGGCGTCGCGCGGAGTTTTCGGGCGTTTCGTTTTTCGGCGCGAGCGGTTTCGGTTCGGGTTCGGCGCGGGGATGGACGGTTTCGGGGTTAGCGTCGTGGTCGTGCGCGTCGGTATTTCGGATTTGTAGTAGA

BC-unmethylated Row 4 :GGTGGGGGGGAAGGGGGGGTTGGGGTGTTGTGTGGAGTTTTTGGGTGTTTTGTTTTTTGGTGTGAGTGGTTTTGGTTTGGGTTTGGTGTGGGGATGGATGGTTTTGGGGTTAGTGTTGTGGTTGTGTGTGTTGGTATTTTGGATTTGTAGTAGA

RefSeq Row 5 : CGgtgagcccCGcCGccctgggccCGgcCGtgcccctgCGctcccCGccCGggattcccccaccccCGcCGggCGCGccCGgCGccCGggaccccCGgcccaCGgctactcacccctccccCGcCGcctcCGcCGggaccctcccc

BC-methylated Row 5 : CGGTGAGTTTCGTCGTTTTGGGTTCGGTCGTGTTTTTGCGTTTTTCGTTCGGGATTTTTTTATTTTCGTCGGGCGCGTTCGGCGTTCGGGATTTTCGGTTTACGGTTATTTATTTTTTTTCGTCGTTTTCGTCGGGATTTTTTTT

BC-unmethylated Row 5 :TGGTGAGTTTTGTTGTTTTGGGTTTGGTTGTGTTTTTGTGTTTTTTGTTTGGGATTTTTTTATTTTTGTTGGGTGTGTTTGGTGTTTGGGATTTTTGGTTTATGGTTATTTATTTTTTTTTGTTGTTTTTGTTGGGATTTTTTTT

RefSeq Row 6 : atCGccagggCGgggcccCGggaagccCGgcccctggggCGgggcttCGgcCGCGgttCGCGgaggCGCGgggtccCGggCGcCGgcaccCGagcccCGgactccttCGgCGggggccCGgggctCGgcacccCGcatggggcCGgCGgg

BC-methylated Row 6 : ATCGTTAGGGCGGGGTTTCGGGAAGTTCGGTTTTTGGGGCGGGGTTTCGGTCGCGGTTCGCGGAGGCGCGGGGTTTCGGGCGTCGGTATTCGAGTTTCGGATTTTTTCGGCGGGGGTTCGGGGTTCGGTATTTCGTATGGGGTCGGCGGG

BC-unmethylated Row 6 :ATTGTTAGGGTGGGGTTTTGGGAAGTTTGGTTTTTGGGGTGGGGTTTTGGTTGTGGTTTGTGGAGGTGTGGGGTTTTGGGTGTTGGTATTTGAGTTTTGGATTTTTTTGGTGGGGGTTTGGGGTTTGGTATTTTGTATGGGGTTGGTGGG

RefSeq Row 7 : gCGggtcCGCGctccCGggacctgagctcaCGagccCGctcCGctgcagaagtgcctgCGcctggaccCGgcCGCGccCGtgtgggcCGccaagcagCGCGtgctctgCGccctcaaccacagcctccaggaCGCGctcaactatgggcttttccagc

BC-methylated Row 7 : GCGGGTTCGCGTTTTCGGGATTTGAGTTTACGAGTTCGTTTCGTTGTAGAAGTGTTTGCGTTTGGATTCGGTCGCGTTCGTGTGGGTCGTTAAGTAGCGCGTGTTTTGCGTTTTTAATTATAGTTTTTAGGACGCGTTTAATTATGGGTTTTTTTAGT

BC-unmethylated Row 7 :GTGGGTTTGTGTTTTTGGGATTTGAGTTTATGAGTTTGTTTTGTTGTAGAAGTGTTTGTGTTTGGATTTGGTTGTGTTTGTGTGGGTTGTTAAGTAGTGTGTGTTTTGTGTTTTTAATTATAGTTTTTAGGATGTGTTTAATTATGGGTTTTTTTAGT

> Supplementary Figure S2

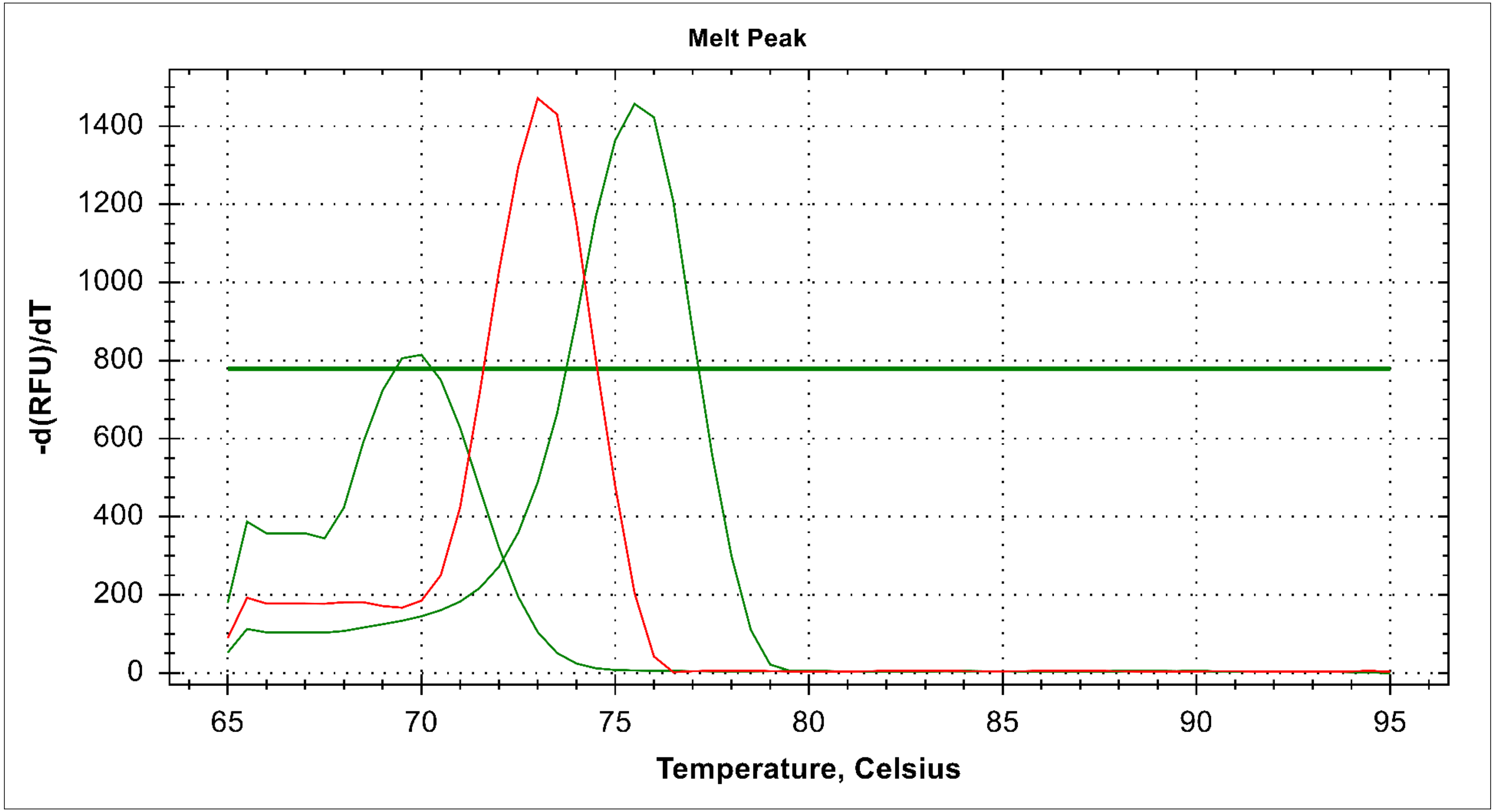

> Supplementary Figure S3

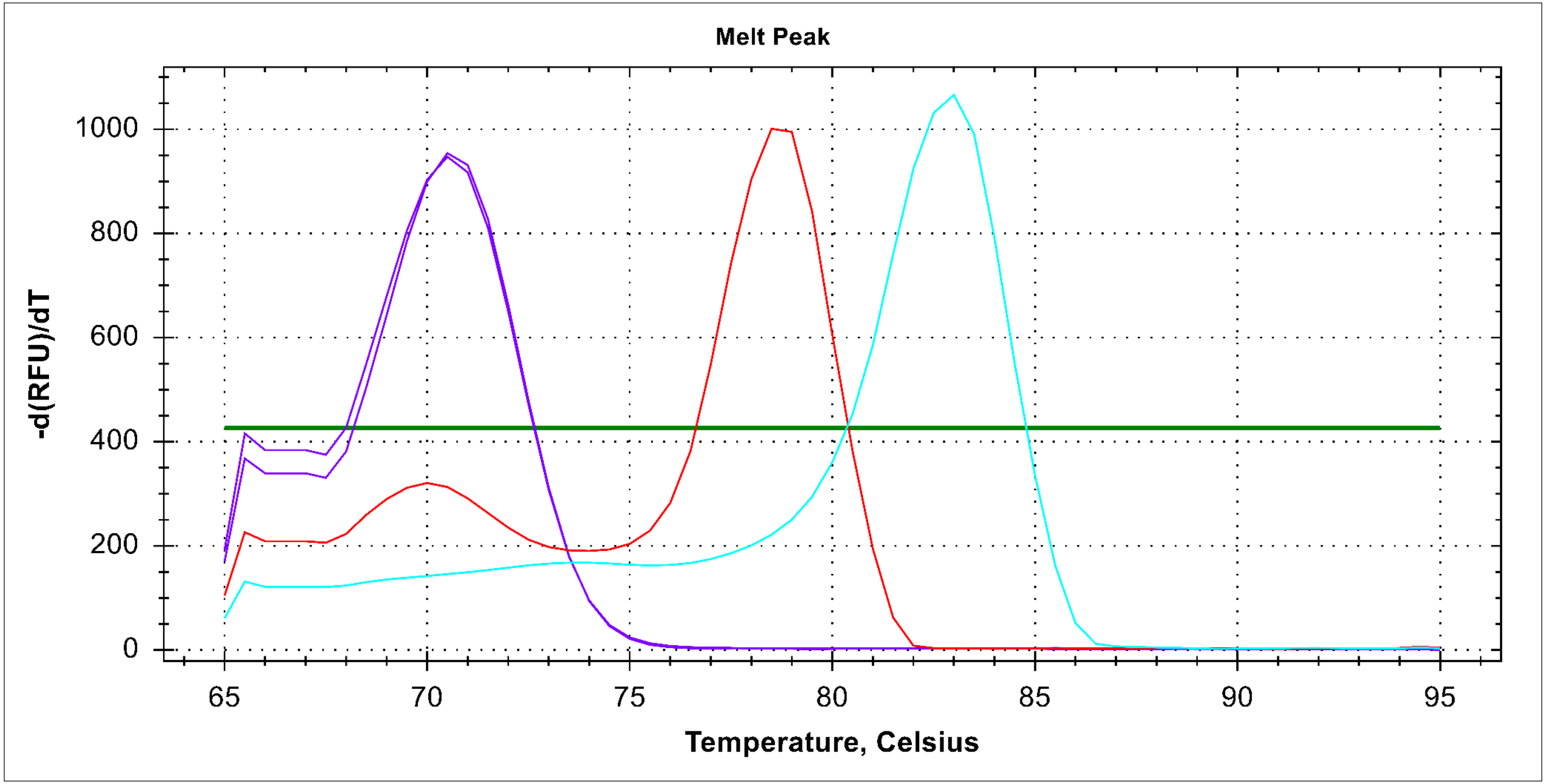
